# *Grin2a* Dysfunction Impairs Cognitive Flexibility by Disrupting LC Modulation of mPFC Circuits

**DOI:** 10.1101/2025.02.01.636062

**Authors:** Hassan Hosseini, Sky Evans-Martin, Emma Bogomilsky, Kevin S. Jones

## Abstract

Cognitive flexibility, a key executive function, is impaired in psychiatric disorders involving prefrontal cortical dysfunction. The medial prefrontal cortex (mPFC) regulates cognitive flexibility and receives noradrenergic input from the locus coeruleus (LC). Mutations in GRIN2A, encoding GluN2A-containing NMDA receptors, impair cognitive flexibility and psychiatric resilience, yet the circuit mechanisms remain unclear. Optogenetic LC→mPFC activation improved reversal learning in wild-type and *Grin2a* heterozygous (HET) mice but not in knockouts (KO), indicating a loss of noradrenergic modulation. *Grin2a* mutants displayed disrupted gamma and high-frequency oscillations (HFOs) in the mPFC. Exogenous norepinephrine failed to restore oscillatory activity, implicating α2-adrenoceptors in NE-driven cortical dynamics. Increased LC innervation and norepinephrine transporter (NET) expression in *Grin2a* mutants suggest excessive noradrenergic input and impaired NE clearance. These findings identify GluN2A as essential for LC-driven prefrontal network synchronization and cognitive flexibility, offering insights into NE dysfunction in psychiatric disorders.

## Introduction

Cognitive flexibility—the ability to adapt behavior in response to changing task demands—is a fundamental component of adaptive functioning and executive control ^1,2^. Deficits in cognitive flexibility are core features of various neuropsychiatric disorders, including schizophrenia, attention-deficit hyperactivity disorder (ADHD), and major depressive disorder, and are associated with poor treatment response and functional impairment ^3–5^. Among cognitive flexibility tasks, reversal learning is particularly sensitive to disruptions in synaptic and neuromodulatory systems ^6,7^.

N-methyl-D-aspartate receptors (NMDARs) play a critical role in regulating cognitive flexibility, with reversal learning showing selective vulnerability to NMDAR dysfunction across spatial and non-spatial tasks ^8–10^. Mutations in GRIN2A, which encode the GluN2A subunit of the NMDAR, are strongly linked to intellectual disabilities, epilepsy, and neurodevelopmental disorders ^11^. Despite this genetic link, the precise mechanisms by which GluN2A dysfunction alters cognitive flexibility remain unclear. Studies in *Grin2a* mutant mice indicate deficits in reversal learning and disrupted network activity, suggesting that GluN2A-containing NMDARs play a crucial role in adaptive behavior ^12–14^.

GluN2A-containing NMDARs are essential for maintaining cortical network stability, particularly by regulating gamma oscillations and synaptic plasticity in the prefrontal cortex (PFC) ^15,16^. Given that cognitive flexibility depends on prefrontal network coordination, GluN2A dysfunction may impair the ability of cortical circuits to adaptively regulate behavior ^17^. While the midbrain dopaminergic system is well known for modulating cognitive processes, the locus coeruleus (LC)—the brain’s primary source of norepinephrine (NE)—provides widespread innervation of the PFC and is essential for reversal learning ^18–20^.

Recent evidence suggests that LC→mPFC circuits dynamically regulate cognitive flexibility ^21–23^. However, it remains unclear how GluN2A dysfunction impacts LC-driven modulation of prefrontal networks. We hypothesize that *Grin2a* loss alters LC-NE signaling by increasing NET expression, reducing extracellular NE, and impairing gamma oscillatory activity, leading to deficits in reversal learning. Using optogenetics, pharmacology, and electrophysiology, we investigate how GluN2A dysfunction disrupts LC→mPFC circuits, highlighting potential therapeutic targets for restoring cognitive function in psychiatric disorders.

## RESULTS

### Reversal Learning is impaired in Grin2a Mutant Mice

While cognitive flexibility is impaired in *Grin2a* mutant mice, previous studies have reported conflicting results regarding their performance in reversal learning tasks. Ryan et al. ^24^ observed significant deficits in reversal learning, whereas Marquardt et al. ^13^ did not identify impairments, raising questions about task-specific or methodological factors influencing these outcomes. To address this discrepancy, we employed a T-maze reversal learning task, which integrates associative and spatial learning processes ^25,26^. Consistent with Ryan et al., we found that *Grin2a*^HET^ and *Grin2a*^KO^ mice required significantly more sessions to reach the criterion compared to WT mice during the reversal phase (**Fig. 1A-B**). These results confirm impaired spatial reversal learning in *Grin2a* mutants and suggest broader deficits in adaptive behavior.

**Figure 1.**
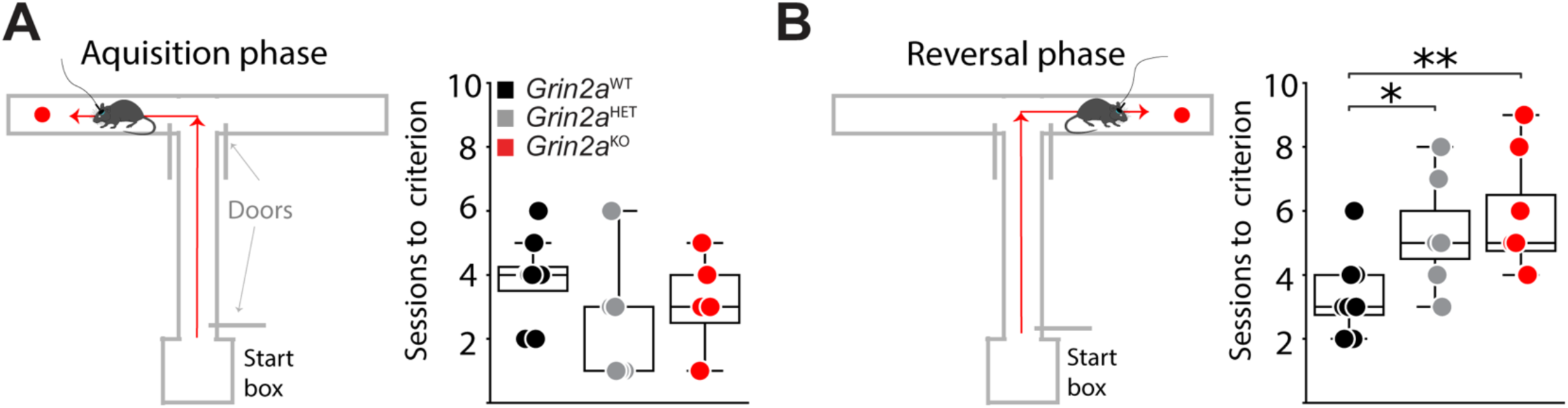
Reversal Learning Is Impaired in *Grin2a* Deficient Mice. **(A)** T-maze acquisition phase schematic. Mice made choice runs toward a sucrose pellet reward (red circle) while controllable doors guided the available options. Box plots represent the number of sessions required to reach criterion (>70% correct responses) for each genotype (WT: 3.88 ± 0.48, *Grin2a*^HET^: 2.29 ± 0.89, *Grin2a*^KO^: 3.00 ± 0.50). **(B)** T-maze reversal learning phase schematic. The reward location was switched, requiring mice to learn the new contingency. Box plots show the number of sessions required to reach criterion (>70% correct responses) in each genotype (WT: 3.38 ± 0.46, *Grin2a*^HET^: 5.29 ± 0.72, *Grin2a*^KO^: 5.75 ± 0.65). Statistical analysis: Kruskal–Wallis test, *n* = 7–8 mice/genotype. Asterisks denote significance: \**p* < 0.05, \*\**p* < 0.01. Box plots: center lines = medians, box edges = 25th and 75th percentiles, whiskers = min and max values.

### LC→mPFC Projections Bidirectionally Modulate Reversal Learning in WT but Not Grin2a Mutants

LC→mPFC projections modulate prefrontal activity through noradrenergic signaling, critical for adaptive behaviors such as cognitive flexibility and reversal learning ^7,25^. Although chemogenetic studies have implicated these circuits in enhancing cognitive flexibility ^23,26,27^, the specific contribution of LC→mPFC activity to reversal learning remains unclear. To directly examine this relationship, we used optogenetics to bidirectionally manipulate LC→mPFC projections during T-maze reversal learning (**Fig. 2A-D**).

**Figure 2.**
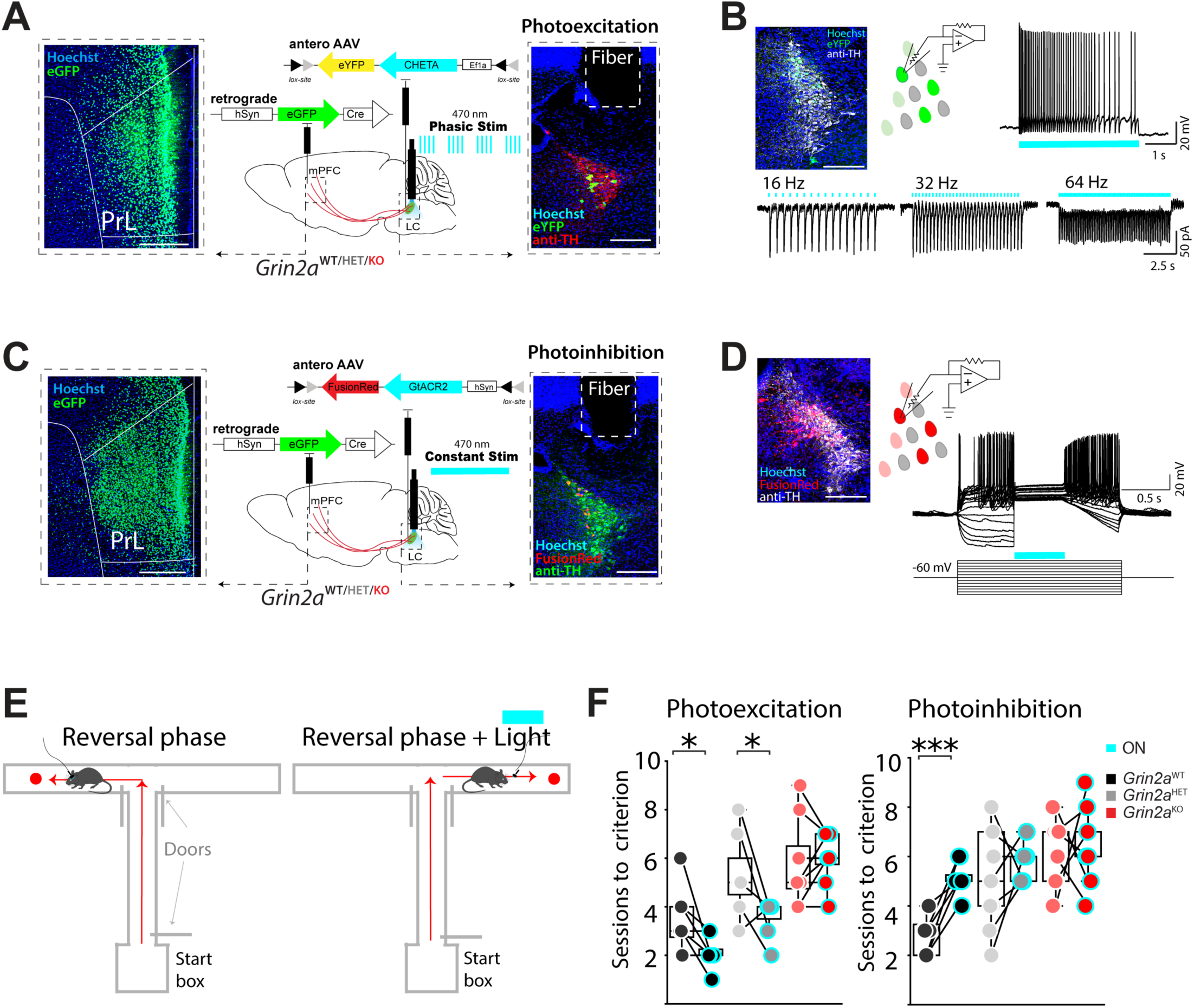
LC→mPFC Projections Bidirectionally Modulate Reversal Learning in WT but Not *Grin2a* Mutants. **(A)** Confocal micrographs and optogenetic schematic. Left: Confocal image showing retro-Cre labeling of mPFC neurons. Right: Optical fiber implantation in the LC for optogenetic stimulation of mPFC-projecting LC neurons. Scale bars: 500 μm (mPFC), 100 μm (LC). Phasic LED stimulation (4 bursts at 25 Hz for 10 s, 470 nm). **(B)** Confocal micrograph of LC section with TH+-stained (white) and retro-Cre labeled neurons that project to the mPFC (green). Voltage-clamp recording from eYFP+ LC neurons. Action potential firing induced by 5 s of 470-nm light (blue bars) at increasing stimulation frequency. **(C)** Confocal micrograph and optogenetic schematic. Same as (A) but with constant 470-nm LED stimulation (10 s total). **(D)** Confocal micrograph of LC section with TH+-stained (white) and FusionRed+ (GtACR2-expressing) LC neurons. Current-clamp recordings from FusionRed+ LC neurons. A 500-ms light pulse (blue bars) suppresses neuronal spiking firing during 1.5- s depolarization steps. **(E)** T-maze reversal learning schematic. Cognitive flexibility was assessed before and after optogenetic stimulation (blue bar). **(F)** Optogenetic modulation of reversal learning. Box plots show sessions required to reach criterion (>70% correct responses) in T-maze reversal: Phasic excitation (470 nm, bursts) improved reversal learning in WT (2.12 ± 0.23) and *Grin2a*^HET^ (3.57 ± 0.31) but had no effect in *Grin2a*^KO^ (6.00 ± 0.38). Constant stimulation (470 nm, sustained) impaired reversal learning in WT (5.12 ± 0.28) but had no effect in *Grin2a* mutants (*Grin2a*^HET^: 5.67 ± 0.24, *Grin2a*^KO^: 6.44 ± 0.50). Statistical analysis: Paired t-test, n = 7– 9 mice/genotype. Asterisks denote significance: \**p* < 0.05, \*\*\**p* < 0.001. Box plots: center lines = medians, box edges = 25th and 75th percentiles, whiskers = min and max values.

Optogenetic activation of the LC→mPFC circuit significantly improved reversal learning performance in WT and *Grin2a*^HET^ mice (**Fig. 2E-F**) but failed to improve performance in *Grin2a*^KO^ mice. Conversely, optogenetic inhibition increased the time required to reach the criterion in WT mice (*p* = 0.0008; **Fig. 2E-F**) but did not affect *Grin2a*^HET^ or *Grin2a*^KO^ mice. These findings demonstrate that LC→mPFC circuits bidirectionally regulate reversal learning in WT mice and highlight the disruption of this mechanism in Grin*2a* mice.

### Enhanced Cortical Projections and Membrane Depolarization of LC Neurons in Grin2a Mice

Excessive noradrenergic (NE) activity in the mPFC, marked by increased innervation and elevated norepinephrine transporter (NET) expression, has been implicated in cognitive flexibility deficits and associated psychiatric conditions ^26,28–30^. To investigate how reduced GluN2A function alters the LC→mPFC circuit, we examined structural and functional changes in *Grin2a* mutants using immunohistochemistry and electrophysiology.

Immunohistochemical analysis revealed a significant ∼2-fold increase in LC innervation density within the PrL of *Grin2a*^HET^ and *Grin2a*^KO^ mice compared to WT (**Fig. 3A-D**).

**Figure 3.**
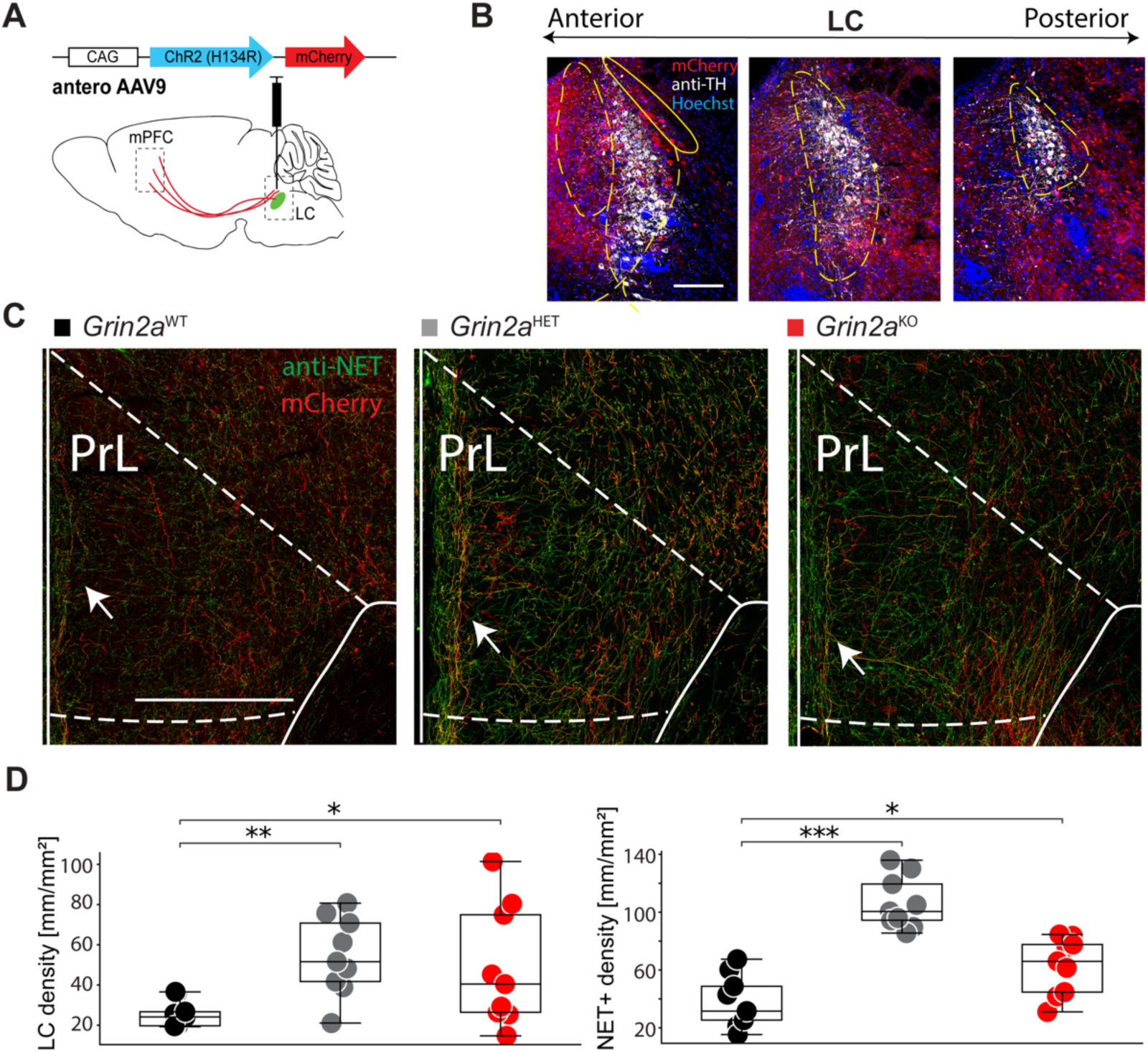
*Grin2a* Mice Exhibit Enhanced LC→mPFC Cortical Projections. **(A)** Viral injection strategy schematic. AAV-mCherry was injected into the LC to label LC neurons. **(B)** Confocal micrographs of TH+ and mCherry+ neurons in anterior and posterior sections of LC. Scale bar: 200 μm. **(C)** LC→mPFC projections in the prelimbic (PrL) region. Confocal micrographs display mCherry+ LC fibers in the PrL of WT, *Grin2a*^HET^, or *Grin2a*^KO^ mice. Scale bar: 500 μm. **(D)** Quantification of LC fiber density. Box plots show fiber density (mm/mm^2^) of mCherry+ LC fibers: WT (25.72 ± 2.20), *Grin2a*^HET^ (54.43 ± 6.48), *Grin2a*^KO^ (48.67 ± 9.96). NET+ fibers: WT (38.13 ± 5.96), *Grin2a*^HET^ (106.31 ± 6.02), *Grin2a*^KO^ (62.96 ± 6.54). Sample size: Mean from three coronal sections (60 μm thick) per mouse, *n* = 3 mice/genotype. Statistical analysis: Kruskal–Wallis test. Asterisks denote significance: \**p* < 0.05, \*\**p* < 0.01, \*\*\**p* < 0.001. Box plots: center lines = medians, box edges = 25th and 75th percentiles, whiskers = min and max values.

Similarly, NET+ axon density was ∼3-fold higher in *Grin2a*^HET^ mice and ∼1.5-fold higher in *Grin2a*^KO^ mice (**Fig. 3A-D**), indicating excessive noradrenergic input to the mPFC.

Electrophysiological recordings revealed further dysfunction in LC neurons of *Grin2a* mutants. The resting membrane potential of LC neurons was depolarized by ∼4.5 mV compared to WT neurons (**Fig. S1A-E**). Despite this, most synaptic properties remained unchanged (**Fig. S2A-D**), with the notable exception of a significant increase in sIPSC frequency observed in LC neurons from *Grin2a*^HET^ mice (p = 0.0025; **Fig. S2B**).

The GluN2A subunit confers rapid decay kinetics to NMDAR current ^31^. Consistent with this role, NMDA currents in LC neurons from *Grin2a*^HET^ and *Grin2a*^KO^ mice decayed significantly more slowly than WT (**Fig. S2A-E**). Furthermore, NMDAR current amplitude was significantly reduced in *Grin2a*^KO^ mice (**Fig. S2E-F**). Collectively, the structural and functional alterations in the LC–mPFC circuit may contribute to deficits in cognitive flexibility observed in *Grin2a* mice.

### Optogenetic Stimulation of LC Terminals Enhances Oscillations in the mPFC of WT but Not Grin2a Mice

Gamma oscillations in the mPFC are essential for cognitive flexibility, acting as regulators of prefrontal processing and adaptive behavior ^32,33^. NE released from LC terminals drives gamma oscillations via volume transmission, modulating cortical network dynamics over extended timescales ^34,35^. Due to the temporal and spatial characteristics of these oscillations, MEA recordings are particularly well-suited for capturing network-wide NE-induced activity.

To investigate how *Grin2a* dysfunction affects LC-NE-induced mPFC oscillations, we conducted MEA recordings in mPFC slices while optogenetically activating LC terminals to mimic endogenous NE release (**Fig. 4A**). In WT slices, optogenetic stimulation caused a 1.5- to 2-fold increase in high-frequency oscillations (iHFO) and gamma-band oscillations (iGBO) in PrL layers 2/3 and 5 (**Fig. 4A**). In contrast, the same optogenetic activation failed to increase oscillatory power in mPFC slices from *Grin2a*^HET^ and *Grin2a*^KO^ (**Fig. 4B-E**). These findings demonstrate that LC-NE signaling is sufficient to drive gamma oscillations in the WT PrL but is impaired in *Grin2a* mutants. This disruption in mPFC network dynamics may contribute to the deficits in cognitive flexibility observed in *Grin2a* mice.

**Figure 4.**
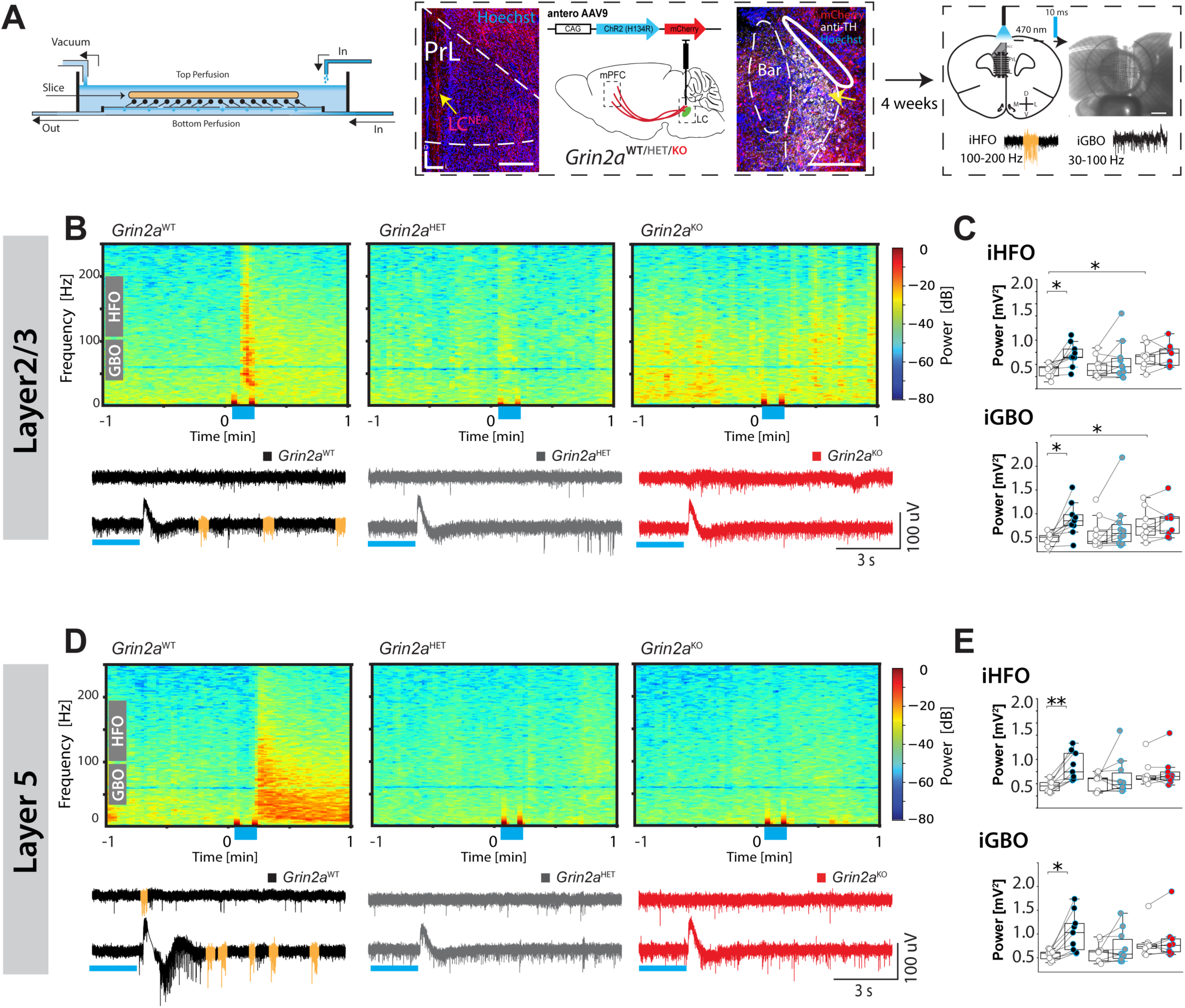
Optogenetic Stimulation of LC Terminals Enhances Oscillations in the mPFC of WT but Not *Grin2a* Mice. **(A)** Left: Schematic of the perfusion-based pMEA chamber with independent top and bottom perfusion routes for the mPFC slice. Middle: Coronal confocal micrograph showing ChR2-expressing, mCherry+ LC terminal projections (red) in the PrL region of the mPFC. Scale bar, 100 μm. Schematic depicts the injection of a ChR2-expressing virus into the LC. Right: Schematic of optogenetic stimulation of LC→mPFC terminals in an *ex vivo* mPFC slice mounted on a pMEA chip and representative traces of optogenetically-induced high-frequency oscillations (iHFO, 100–200 Hz; highlighted in orange) and gamma-band oscillations (iGBO, 30–100 Hz). Scale bar, 1 mm. (**B**) Representative spectrograms and local field potential (LFP) recordings (HFO events highlighted as orange) from layer II/III of the PrL following optogenetic activation (blue box) of LC terminals in WT and *Grin2a* mutant mice. (**C**) Box plots of iHFO and iGBO power in layer II/III (baseline vs. post-stimulation). (**D**) Representative spectrograms and LFP recordings from layer V PrL of WT and *Grin2a* mutant mice (same conditions as in **B**). (**E**) Box plots of iHFO and iGBO power in layer V (baseline vs. post-stimulation). Data in (**C, E**) were analyzed via a two-way ANOVA with Tukey’s post-hoc test for multiple comparisons. *n* = 8–9 slices from 3 mice/genotype. See Table 1 for analyses. Asterisks denote significance: \**p* < 0.05, \*\**p* < 0.01. Box plot centers represent medians, box edges the 25th and 75th percentiles, and whiskers the minimum and maximum data points.

### Exogenous NE Induces Oscillations in WT mPFC but Not in *Grin2a* Mutants

The inability of optogenetic activation of LC terminals to induce oscillations in *Grin2a* mutants suggests a disruption in noradrenergic signaling, potentially arising from pre-synaptic dysfunction, post-synaptic dysfunction, or a combination of both. To distinguish between these possibilities, we assessed oscillatory responses to exogenous NE application, bypassing the LC terminals (**Fig. 5A**).

**Figure 5.**
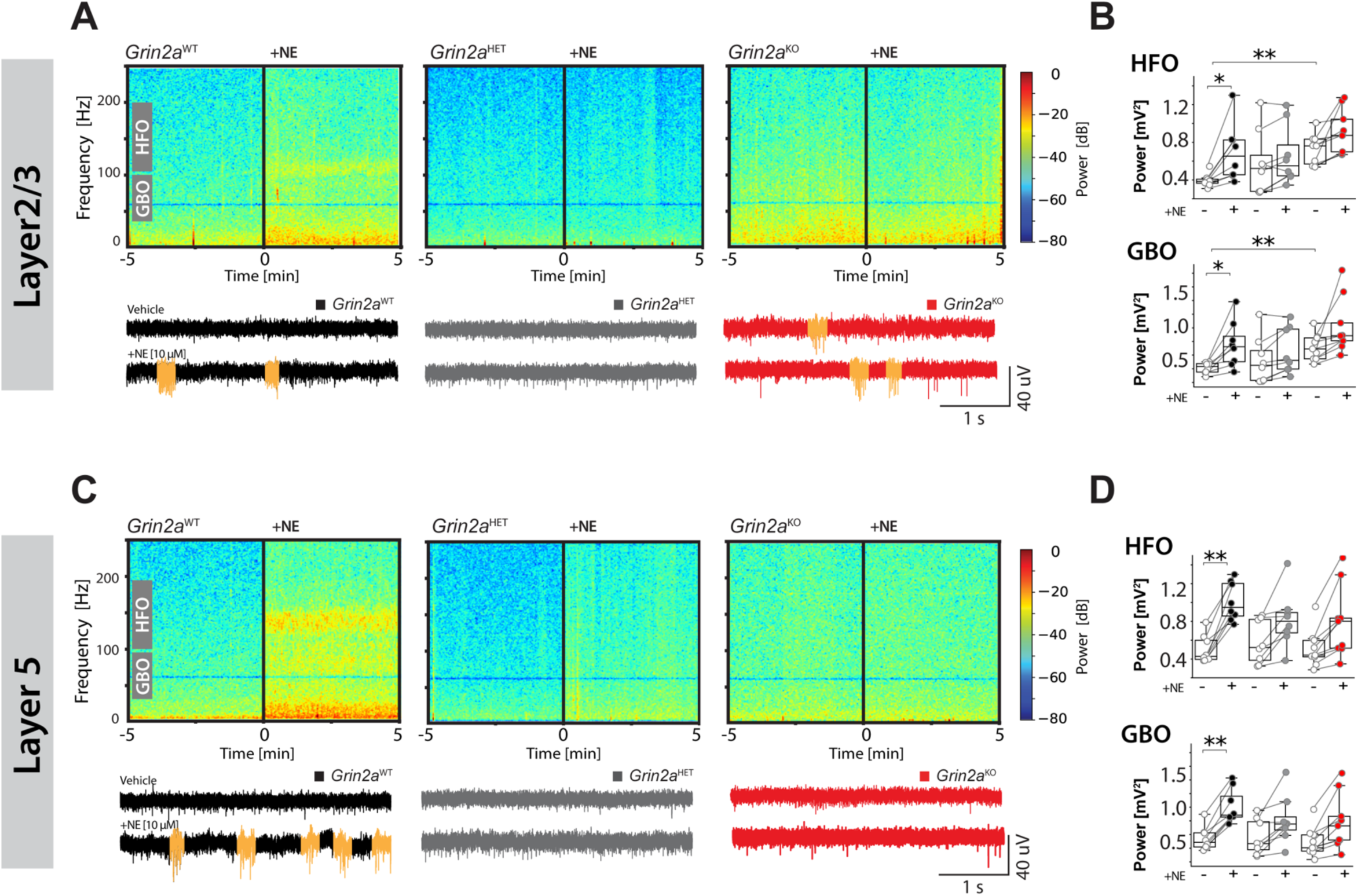
Exogenous NE Induces Oscillations in WT mPFC but Not in *Grin2a* Mutants. (**A, C**). Representative spectrograms and LFP recordings during bath application of norepinephrine (NE, 10 μM) from PrL layer II/III (**A**) or layer V (**C**) from WT, *Grin2a*^HET^, and *Grin2a*^KO^ slices. HFO events highlighted in yellow. (**B, D**) NE-evoked changes in HFO and GBO power in layer II/III (**B**) or layer V (**D**) of PrL slices from WT, *Grin2a*^HET^, and *Grin2a*^KO^ slices. Box plots compare baseline vs. post-NE stimulation. Statistical analysis: Two-way ANOVA with Tukey’s post-hoc test, *n* = 8–9 slices from 3 mice per genotype. Asterisks denote significance: \**p* < 0.05, \*\**p* < 0.01. Box plots: center lines = medians, box edges = 25th and 75th percentiles, whiskers = min and max values.

Basal GBO power in layer 2/3 of the PrL was ∼2-fold higher in *Grin2a*^KO^ slices than in WT and *Grin2a*^HET^ slices (*p* = 0.035; **Fig. 5B-C**). In WT slices, bath-applied NE robustly increased both GBO and HFO power in layers 2/3 and 5 (*p* < 0.001; **Fig. 5B-E**).

However, NE failed to increase HFO or GBO power in *Grin2a*^HET^ or *Grin2a*^KO^ slices (**Fig. 5B-E**). These findings indicate that post-synaptic dysfunction, rather than pre-synaptic mechanisms, is the primary driver of impaired noradrenergic signaling in the mPFC of *Grin2a* mutants.

### Acute GluN2A Blockade Fails to Replicate Oscillatory Deficits in *Grin2a* Mutants

To determine whether the NE-induced oscillatory deficits in *Grin2a* mutants result from acute disruptions in GluN2A signaling or developmental alterations, we applied NVP, a selective GluN2A antagonist, to mPFC slices from WT mice. This approach allowed us to assess whether acute GluN2A blockade replicates the oscillatory impairments observed in *Grin2a* mutants. As expected, bath-applied NE increased HFO and GBO power in the WT PrL by approximately 2-fold (GBO: *p* = 0.018; HFO: *p* = 0.0046; **Fig. 6A-B**), establishing a baseline for evaluating the role of GluN2A. Notably, concurrent application of NVP did not significantly alter GBO or HFO power compared to NE alone (*p* = 1.00; **Fig. 6A-B**). These results indicate that acute GluN2A signaling is not essential for NE-induced oscillations in the mPFC. Instead, the absence of NE-induced oscillatory activity in *Grin2a* mutants likely reflects disruptions in the developmental or maturational processes that shape prefrontal circuitry.

**Figure 6.**
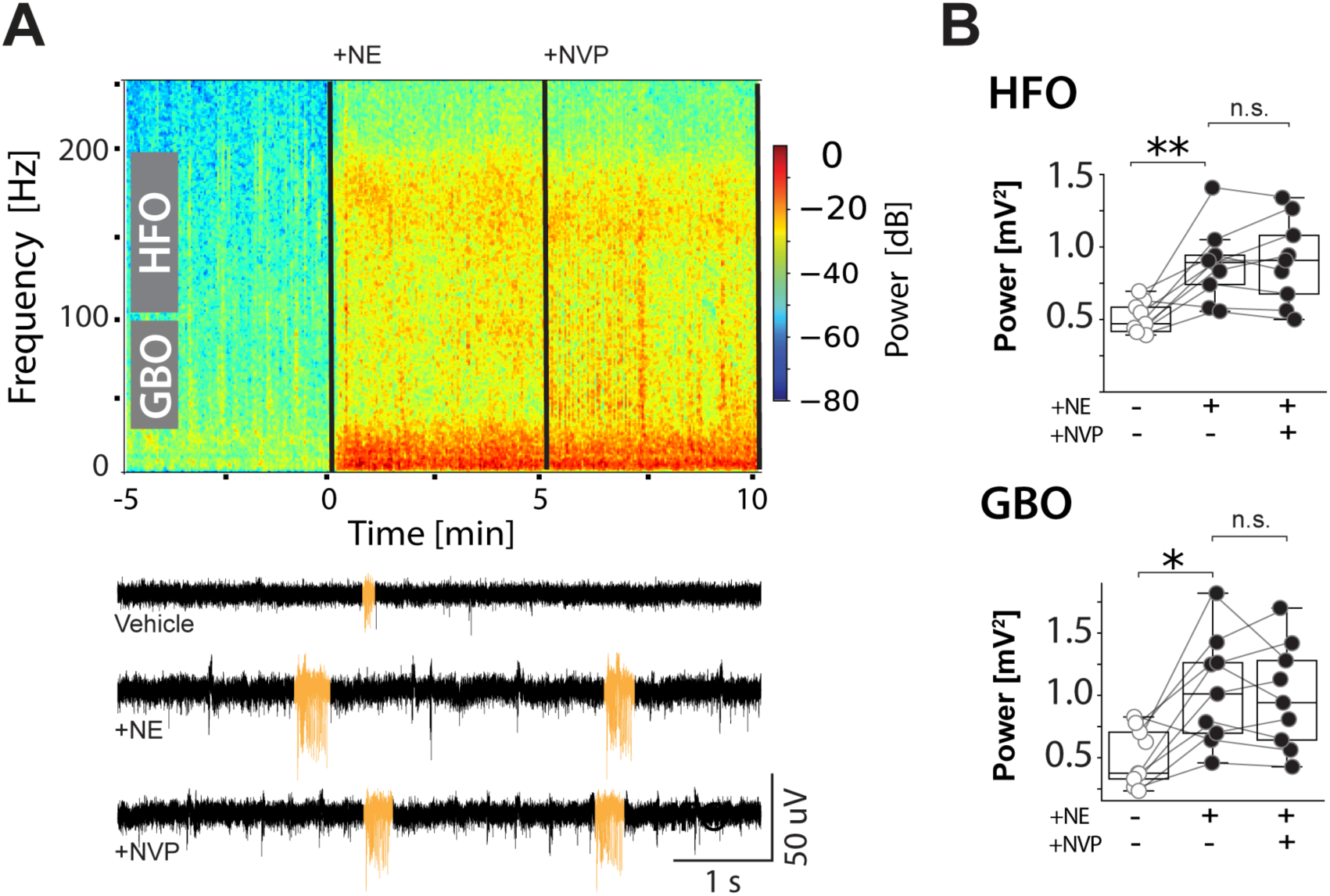
Acute GluN2A Blockade Fails to Replicate Oscillatory Deficits in *Grin2a* Mutants. (**A**) Representative spectrogram and LFP recording (beow) from an electrode in layer 5 of WT PrL illustrating oscillatory responses to norepinephrine (NE, 10 μM) alone or with the GluN2A antagonist NVP-AAM077 (0.5 μM). (**B**) Quantification of HFO and GBO power. Box plots compare baseline (vehicle), NE (10 μM), and NE + NVP (0.5 μM) conditions: HFO power (mV^2^): Vehicle (0.409 ± 0.029), NE (0.703 ± 0.069), NE + NVP (0.721 ± 0.078). GBO power (mV^2^): Vehicle (0.150 ± 0.015), NE (0.258 ± 0.029), NE + NVP (0.248 ± 0.028). Statistical analysis: Kruskal– Wallis test, n = 9 slices from 3 WT mice. Asterisks denote significance: *p < 0.05, **p < 0.01. Box plots: center lines = medians, box edges = 25th and 75th percentiles, whiskers = min and max values.

### α2 Adrenergic Receptors Are Essential for NE-Induced mPFC Oscillations

Having determined that acute GluN2A signaling is not required for NE-induced oscillations, we next explored the role of adrenergic receptors, which regulate mPFC network dynamics and adaptive behaviors ^27,35^. Given the established role of α2 adrenergic receptors in modulating PFC function ^36–38^, we investigated their involvement in NE-induced oscillations.

Bath application of the α2-selective antagonist, yohimbine, abolished NE-induced increases in HFO and GBO power in WT mPFC slices (GBO: *p* = 0.016; HFO: *p* = 0.025; **Fig. 7B-C**). In contrast, selective antagonists for α1, β1, and β2 adrenergic receptors had no significant effect on NE-induced oscillations (**Fig. 7B-C**). Furthermore, yohimbine eliminated oscillatory activity generated by optogenetic activation of LC terminals in the mPFC (**Table 1-2**). These findings establish α2 adrenergic receptors as essential mediators of mPFC oscillations driven by both endogenous and exogenous NE. The disruption of NE-induced oscillations in the mPFC of *Grin2a* mutants underscores the reliance of cortical network stability on α2 signaling and suggests that GluN2A deficiency may impair α2 receptor-mediated modulation of network activity, potentially through disrupted developmental or synaptic mechanisms.

**Figure 7.**
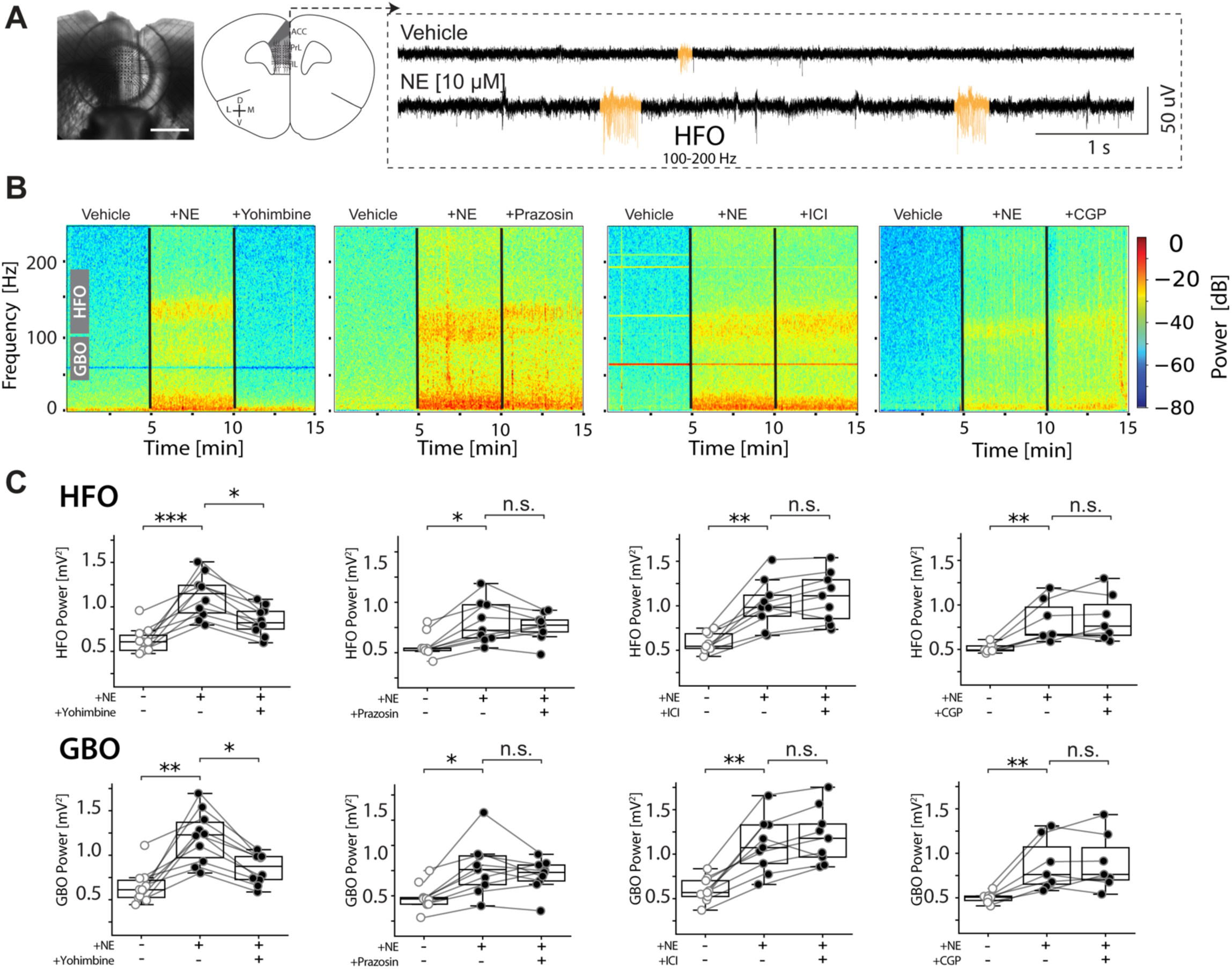
α2 Adrenergic Receptors Are Essential for NE-Induced mPFC Oscillations. **(A)** Left: Bright-field micrograph of *ex vivo* mPFC slice on pMEA chip (scale bar: 1 mm). Middle: Schematic showing region of interest. Right: Prototypical LFP trace from layer 5 PrL electrode in vehicle or norepinephrine (NE, 10 μM). HFO events highlighted in yellow. **(B)** Representative spectrograms from a L5 PrL electrode in response to NE and subsequent co-treatment with a subtype-selective adrenergic receptor antagonists: Yohimbine (α2, 10 μM); Prazosin (α1, 100 nM); ICI-118551 (β2, 10 nM); CGP-20712A (β1, 100 nM). **(C)** Quantification of pharmacologically modulated changes in HFO and GBO power. Plots compare mean oscillatory power per treatment. Statistical analysis: Two-way ANOVA with Tukey’s post-hoc test for multiple comparisons, *n* = 8–9 slices from 3 WT mice. Asterisks denote significance: \**p* < 0.05, \*\**p* < 0.01, \*\*\**p* < 0.001. Box plots: center lines = medians, box edges = 25th and 75th percentiles, whiskers = min and max values.

## DISCUSSION

In this study, we demonstrate that GluN2A-containing NMDARs are pivotal for regulating both reversal learning and cortical oscillations via noradrenergic signaling in the mPFC. We found that *Grin2a* ablation impairs reversal learning and disrupts both noradrenergic input and network dynamics in the mPFC. To our knowledge, this is the first causal evidence showing that optogenetic stimulation or inhibition of the LC→mPFC circuit can enhance or diminish spatial reversal learning. Furthermore, the structural and functional abnormalities identified here suggest that alterations in LC→mPFC circuitry are key contributors to the cognitive deficits observed in *Grin2a* mice, with implications for the pathophysiology of neuropsychiatric disorders.

### LC→ mPFC Modulation of Reversal Learning Requires Developmental GluN2A Function

Building on these initial observations, our findings reveal that LC→mPFC circuits are critical for spatial reversal learning. Optogenetic activation of LC→mPFC projections improved reversal learning in WT and *Grin2a*^HET^ mice but not in *Grin2a*^KO^ mice (**Fig. 2**), while inhibition impaired performance only in WT mice. This contrast highlights the role of GluN2A signaling in enabling noradrenergic modulation of cognitive flexibility.

Reversal learning, like many forms of cognitive flexibility, improves with development and peaks in early adulthood ^39,40^. During adolescence, NET expression in the PrL is elevated and then declines in adulthood ^41^. Environmental stressors and elevated NET expression impair cognitive flexibility, whereas pharmacological blockade of NET can restore flexibility via enhanced noradrenergic signaling ^41^.

In *Grin2a* mutants, a 1.5–3-fold increase in NET expression (**Fig. 3**) suggests excessive NE reuptake in the PrL, thereby reducing extracellular NE levels and disrupting its modulatory impact on gamma oscillations. Together, these findings indicate that cognitive deficits in *Grin2a* mutants arise from impaired noradrenergic modulation, driven by excessive NE reuptake and disrupted LC physiology. Furthermore, GluN2A signaling is essential for maintaining the functional integrity of the LC→mPFC pathway. While acute GluN2A blockade does not abolish NE-induced oscillations (**Fig. 6**) ^42^, the chronic absence of GluN2A during development disrupts LC→mPFC circuit function, likely contributing to the deficits in adaptive reversal learning seen in *Grin2a* mutants ^13,24^. Such deficits align with the broader literature on cognitive flexibility impairments across multiple neuropsychiatric disorders ^4,43^.

### α2 Adrenoreceptors Mediate NE-Induced Network Oscillations

Our findings indicate that, in *Grin2a* mutants, neither optogenetic stimulation of LC terminals nor exogenous NE application evokes GBO or HFO in layer 5 of the mPFC (**Fig. 4,5**), suggesting impaired LC-NE modulation of cortical oscillations. This disruption in oscillatory activity likely contributes to the reversal learning deficits observed in these mice. NE-induced oscillations in the mPFC are mediated primarily via α2 adrenergic receptor signaling ^36–38^. Moreover, pharmacological blockade of α2 receptors with yohimbine abolished both NE-induced and optogenetically driven increases in GBO and HFO power, whereas selective antagonists of α1, β1, or β2 had no significant effect (**Fig. 7**). This was further supported by a broad suppression of oscillatory power across frequency bands following α2 receptor blockade (**Fig. 7B-C**; **Table 1-2**).

Hence, the reduced efficacy of α2 receptor signaling in *Grin2a* mutants likely contributes to their cognitive flexibility deficits, highlighting α2 receptors as pivotal regulators of cortical network function with potential therapeutic relevance in NMDAR hypofunction.

Impaired gamma oscillatory activity, rooted in disrupted α2 receptor function, may also reflect broader deficits in prefrontal network synchronization—an essential component of adaptive behavior ^44,45^.

### Mechanistic Insights: LC-NE Dysfunction and Cognitive Deficits

Building on prior work demonstrating that GluN2A signaling maintains E/I balance and network stability in PFC circuits ^46^, we now show that *Grin2a* ablation differentially impacts noradrenergic modulation, offering key mechanistic insights into how NMDAR hypofunction disrupts cortical networks. In addition to delaying the maturation of parvalbumin interneurons and disrupting synaptic integration ^47^ ^46^, *Grin2a* ablation also impairs hippocampal oscillatory activity ^48^, implicating widespread network alterations. Extending these findings, the present study identifies deficits in LC-NE circuits, suggesting that noradrenergic dysfunction plays a central role in the behavioral and network-level impairments observed in *Grin2a* mutants.|

Disruptions in gamma oscillatory activity observed in *Grin2a* mutants likely reflect a breakdown in noradrenergic modulation via α2 receptors. Moreover, the depolarized resting potential of LC neurons in *Grin2a* mutants may underlie heightened NE release and enhanced LC→mPFC innervation, further contributing to disrupted cortical dynamics. Such structural and functional abnormalities likely impair cognitive flexibility, a core deficit in psychiatric disorders, including schizophrenia. Beyond the LC-NE axis, our findings also bear relevance to dopaminergic dysfunction. They align with the dopamine hypothesis of schizophrenia, which posits that dysregulated dopaminergic transmission contributes to key cognitive and behavioral symptoms ^49,50^. Given that NE and dopamine (DA) systems are intricately linked—particularly in the PFC, where NE can modulate DA release and receptor sensitivity ^51,52^—disruptions in NE-induced cortical oscillations observed in *Grin2a* mutants could also interfere with DA regulation. This highlights how NMDAR hypofunction might disrupt both NE and DA homeostasis in the PFC, exacerbating core cognitive deficits in psychiatric disorders.

### Alignment with Prior Studies

Our results are consistent with prior research linking *Grin2a* mutations to cognitive and neurodevelopmental deficits^11^. For instance, Kinney et al. ^53^ and Hosseini et al. ^48^ reported disrupted E/I balance and impaired gamma oscillations in hippocampal circuits—findings extended to the mPFC by our previous work ^46^. Additionally, work by Bertocchi et al. and Brigman et al. ^16,17^ demonstrated that GluN2A deficiency impairs reversal learning and cognitive flexibility, findings corroborated by our T-maze results. While earlier studies emphasized the role of GluN2A in hippocampal-dependent learning and memory, our data highlight its critical function in modulating LC→mPFC circuits and cortical oscillatory dynamics, broadening our understanding of how GluN2A dysfunction can affect both network-level coordination and behavior.

Similarly, the gamma oscillation disruptions observed in *Grin2a* mutants echo studies highlighting the necessity of oscillatory synchronization for flexible behavior and cognitive adaptation ^54^. By identifying structural and functional deficits in the LC→mPFC circuit, our findings not only align with prior research on hippocampal circuitry but also suggest that some behavioral phenotypes may arise from hippocampal dysregulation in *Grin2a* mutants. The improvement seen with LC→mPFC stimulation in *Grin2a*^HET^ mice further indicates that bolstering this circuitry may compensate for hippocampal deficits, whereas photoinhibition has little effect—perhaps due to dominant hippocampal dysfunction that cannot be exacerbated further. This more comprehensive understanding of GluN2A dysfunction extends beyond hippocampal pathways to include higher-order cortical areas, revealing how widespread network alterations can underlie cognitive impairments in *Grin2a* mutants.

### Clinical and Translational Implications

The insights gained from our *Grin2a* mouse model highlight potential avenues for therapeutic intervention. Given the clear role of α2 adrenergic receptors in NE-driven oscillations, drugs that boost α2 receptor function could theoretically help normalize prefrontal gamma rhythms and thus mitigate cognitive deficits in disorders with NMDAR hypofunction. Additionally, strategies aimed at stabilizing cortical network dynamics— such as neuromodulation or pharmacological modulation of inhibitory circuits—may complement interventions targeting noradrenergic dysfunction.

Moreover, because chronic GluN2A dysfunction disrupts mPFC maturation, interventions introduced during critical developmental windows may offer the greatest impact on cognitive flexibility and network stability. Finally, the observed interplay between NE and DA systems suggests that combined pharmacological targeting of these pathways may hold promise. Interventions that bolster NE-DA coupling could promote prefrontal stability and improve executive functions disrupted in psychiatric disorders. Future clinical approaches may focus on synergy between pharmacological agents that enhance α2 or DA signaling and neuromodulatory techniques (e.g., transcranial magnetic stimulation), providing a multifaceted strategy for restoring adaptive behavior.

### Future Directions

Our findings underscore the vital role of LC→mPFC circuits in cognitive flexibility amid GluN2A dysfunction, yet several key questions remain. Although NMDAR hypofunction is a central model for psychiatric disorders, it is not the only pathophysiological mechanism at play. Future studies should investigate whether LC→mPFC dysfunction constitutes a convergent deficit across other models—genetic, environmental, or pharmacological—to reveal fundamental processes underlying cognitive deficits.

Additionally, because this circuit also modulates dopaminergic signaling, understanding how GluN2A deficiency affects DA neurotransmission could illuminate the broader network interactions that shape cognitive function. Given the intricate molecular coupling of NE and DA systems—via transporters such as NET and DAT and potential heterodimeric receptors ^52^—further research on NE–DA interactions in *Grin2a* mutants is warranted.

Longitudinal, developmental studies in *Grin2a* mutants will help determine the spatiotemporal windows during which NE and DA circuits most strongly influence mPFC-dependent tasks like reversal learning and set-shifting. Targeting α2 adrenergic receptors or inhibitory interneurons during these intervals could serve as a strategy to prevent or mitigate cognitive impairments associated with NMDAR hypofunction.

While this study focused on spatial reversal learning, examining how LC→mPFC dysfunction impacts a broader range of cognitive domains (e.g., working memory, attention, and decision-making) will further elucidate how these circuits support higher-order cognition. As we broaden the range of behaviors assessed, from working memory to attention and decision-making, uncovering how LC→mPFC dysfunction contributes to diverse psychiatric conditions may open new therapeutic avenues.

## Conclusion

In summary, our findings establish that GluN2A-containing NMDARs are essential for the interplay between NMDAR signaling and LC-NE modulation in the mPFC. We demonstrate, for the first time, that optogenetic manipulation of LC→mPFC activity can both enhance and impair reversal learning, emphasizing the circuit’s pivotal role in adaptive behavior. Further, we show α2 adrenoceptors to be crucial mediators of NE-driven cortical oscillations, suggesting they are prime candidates for therapeutic intervention in conditions involving NMDAR hypofunction.

Taken together, these insights shed new light on how disrupted LC→mPFC communication underlies cognitive deficits in disorders characterized by GluN2A and broader NMDAR dysfunction, paving the way for targeted strategies to restore adaptive behavior.

## METHODS

### Animals

All animal procedures were approved by the University of Michigan’s Institutional Animal Care and Use Committees (IACUC) and conformed to NIH Guidelines for animal use.

*Grin2a^KO^*mice (Riken B6;129S-*Grin2a*; RBRC02256) were intercrossed or outcrossed to C57BL/6J mice to generate *Grin2a*^HET^ or *Grin2a*^KO^, respectively.

Genotypes were confirmed by real-time PCR assay (Transetyx, Cordova, TN). *Grin2a* mutant mice were fostered by WT dams to standardize parental influence on behavior and physiology. Male mice were exclusively used in this study to eliminate variability introduced by the female estrous cycle, ensuring more consistent and reliable results.

### Optogenetic Experiments

#### Surgery

Stereotactic surgeries were performed on 7-8-week-old male mice as previously described ^46^. Briefly, mice were anesthetized and transferred to the stereotactic apparatus. A craniotomy was performed over the target site. For electrophysiology experiments, 110 nl of the anterograde virus pAAV.CAG.hChR2(H134R)- mCherry.WPRE.SV40 (Addgene viral prep #100054-AAV9) was delivered unilaterally into the locus coeruleus (LC, coordinates in mm: AP: −5.4, ML: +0.85, DV: −4.0). For behavioral experiments, retrograde virus pENN.AAV.hSyn.HI.eGFP-Cre.WPRE.SV40 (Addgene viral prep #105540-AAVrg) was unilaterally injected into the right medial prefrontal cortex (mPFC, coordinates in mm: AP: 1.9, ML: +0.3, DV: −1.8) and the LC was injected with either Cre-dependent channelrhodopsin pAAV-Ef1a-DIO-ChETA-EYFP (Addgene viral prep #26968-AAV9) or pAAV-hSyn1-SIO-stGtACR2-FusionRed (Addgene viral prep #105677-AAV1). After viral injections were complete, we implanted a ferrule-coupled optical fiber (0.22 NA, 200-μm diameter, RWD, Catalog No. 807- 00022-00 ) 300 µm above the center of the injection (coordinates in mm: AP: -5.4, ML: +0.85, DV: -3.7). We targeted the LC for unilateral transduction to prevent excessive disruption of general arousal caused by bilateral perturbation ^25^. The fiber implant was fixed by gluing it to the skull with dental cement. After surgeries, mice were administered carprofen (5 mg/kg) to provide analgesia.

#### Optogenetic Intervention

Pathway-specific neural modulation of LC projections to the mPFC was achieved via optogenetic excitation (25-Hz pulses) or inhibition (constant illumination) using 470-nm light (10–17 mW). Light was delivered for 10 seconds through a 200-µm diameter, 0.39 NA fiber optic cable and a fiber optic rotary joint (Thorlabs, RJPSL2). Perturbation occurred during the second run of the reversal learning task, specifically when the mouse was in the start box just before the door opened.

### Electrophysiology

#### Acute slice preparation

Coronal mPFC slices (300 μm thick) were prepared from 12- to 15-week-old male mice as previously described ^48^. In summary, brains were swiftly removed after isoflurane anesthesia and decapitation, submerged in ice-cold, carbogenated slicing solution, sectioned with a vibratome, and subsequently incubated in recovery solution before recordings.

#### Patch-Clamp

Whole-cell patch-clamp recordings were performed from 12- to 15-week-old male mice as previously described ^46^. Briefly, putative LC neurons were characterized visually by their anatomic location on the later side of the fourth ventricle or from retro-grade labeled eYFP+ or FusionRed+ neurons. For NMDAR–mediated postsynaptic currents, neurons were voltage clamped at +40 mV in modified recording ACSF (1 mM MgSO4), and to block α-amino-3-hydroxy-5-methyl-4-isoxazolepropionic acid (AMPA) and gamma-aminobutyric acid (GABA) current, AMPA receptor antagonist CNQX (10 µM) and GABA receptor antagonist picrotoxin (PTX, 10 µM) were added to the ACSF.

#### Multi-electrode Array

Local field potentials (LFPs) were recorded from mPFC coronal slices following previously described methods ^48^. Recordings were conducted after a 15-minute baseline period. Optical stimulation (470 nm, 10 s constant light) was delivered using a digital micromirror device (Polygon-400, Mightex) through a 10x-objective (Olympus). Optical power output was maintained at 1 mW/mm^2^.

#### Histology

Tissue was prepared, permeabilized, and incubated with antibodies as previously described ^46^. Primary antibody: rabbit anti-TH (Millipore, Catalog No. AB152) with 1:1000 dilution and mouse anti-NET (Sigma, Catalog No. AMAb91116) with 1:1000 dilution. Secondary antibodies: Alexa-647 donkey anti-rabbit (Life Technologies #A31573) with 1:2000 dilution and Alexa-488 donkey anti-mouse (Life Technologies #A21202) with 1:2000 dilution. Stained sections (60 μm) were mounted on slides, coverslipped with ProLong Gold Antifade mounting reagent (Molecular Probes, #P36930) and visualized using a confocal microscope. We quantified axonal density by measuring the total axon length in millimeters per square millimeter of tissue using the ’Ridge Detection’ plugin in ImageJ. Native fluorescence was used to detect the mCherry+ signal in LC fibers, and NET+ fibers were labeled via immunohistochemistry to calculate density in the mPFC.

#### Drugs

Norepinephrine hydrochloride (item No. 35580), yohimbine hydrochloride (item No. 19869), prazosin hydrochloride (item No. 15023), ICI-118551 hydrochloride (item No. 15591), CGP-20712A (item No. 40765), NVP-AAM077 (item No. 30622), CNQX (item No. 14618), and picrotoxin (item No. 20771) were purchased from Cayman (Ann Arbor, Michigan, USA). The compounds were dissolved at the specified final concentration in ACSF and added to the bath. To protect the compounds from degradation, solutions containing reagents were freshly prepared before the application. If a compound was dissolved in DMSO, then the same concentration of DMSO was added to the extracellular solution for the control recordings.

#### Behavioral Setup and Training

Behavior training for the reversal learning task started ∼2 weeks after the surgery and is adopted from the T-maze reversal learning task used in a previous study (Maas et al., 2020). Mice were maintained on a food restriction schedule, kept at 75–85% of their free-feeding weight, and food was restricted for 12 hours before each experiment. Mice underwent two sessions (days) of habituation to the T-maze (50 × 7 × 20 cm arms; 15 × 15 cm start box with controllable doors), consisting of 10-minute sessions of free exploration with all doors open and sucrose rewards available. Over the subsequent two sessions, mice underwent 10 trials per day of behavioral shaping, running to baited goal arms in alternating directions for a 1 mg sucrose pellet reward. Mice meeting the speed criterion (1-3 min/trial) proceeded to T-maze reversal training until achieving >70% correct trials per session. During the training phase, mice completed 10 trials per session, selecting a pseudorandomized goal arm (right or left) until they reached a >70% performance criterion. After entering the goal arm, the alternate arm was blocked by a controllable door. In the reversal learning phase, the goal arm was switched, and mice were again tested to achieve >70% accuracy. In subsequent sessions (Reversal phase + optogenetics), the goal arm was switched once more, but this time, mice received optogenetic stimulation—either photo-excitation (25 Hz burst stimulation) or photo-inhibition (10-second constant stimulation)—immediately before leaving the start box. The 25 Hz stimulation frequency was chosen based on previous findings linking LC firing during enhanced memory performance ^55^.

#### Electrophysiological Data Processing and Statistical Analysis

MEA recordings: Spontaneous oscillations were analyzed from 1-min segments before and after optogenetic stimulation of LC terminals in the mPFC. NE-induced oscillations were analyzed from 5-min segments before and after NE bath application in the mPFC. A notch filter was applied to remove 60 Hz noise. The 30–100 Hz range was designated as gamma ^56^ and 100-200 Hz to high-frequency oscillation ^57^. Absolute power within each frequency band was calculated by integrating the power spectrum. Data was processed with custom Python scripts. LFPs were low-pass filtered at 500 Hz and downsampled to 1 kHz. Multi-taper spectral analysis was used for power estimation.

Relative power changes were computed as ((induced − baseline)/baseline). All analysis code is available at https://github.com/NeuroDataa/MEA_Analysis.

Patch-Clamp recordings: Data was analyzed in Python. Passive and active properties were extracted using the eFEL package ^58^, with additional custom Python scripts available at https://github.com/NeuroData/PatchClampAnalysis. For sEPSC, sIPSC and sNMDA detection, traces were filtered at 1 kHz, and events were identified using template matching in pClamp (Clampfit). Templates were defined from averages of >100 events.

## Acknowledgements

This work was supported by the Stanley Center for Psychiatric Research at the Broad Institute (KSJ). The authors report no biomedical financial interests or potential conflicts of interest. We thank Maxwell Kenny for the design and construction of the automated T-Maze apparatus. We thank Dr. Cagney Coomer and Dr. Nandkishore Prakash for critically reviewing the manuscript.

## Author Contributions

K.S.J. conceptualized and designed the study. H.H., S.E.M., and E.B. conducted the study and performed analysis. K.S.J. secured the funding. H.H. developed the methodology, and H.H., S.E.M., and E.B. conducted the experiments. K.S.J. oversaw project administration and supervision. H.H. and K.S.J. wrote the original draft of the manuscript. All authors reviewed and edited the manuscript, contributing to interpreting the results and providing critical feedback throughout the research process.

## Data and Code Availability

All data generated or analyzed during this study are included in this published article (and its supplementary information files) and are available from the sorresponsding author upon reasonable request. Original code has been deposited at https://github.com/NeuroData/PatchClampAnalysis. https://github.com/NeuroDataa/MEA_Analysis.

---

**Figure S1.**
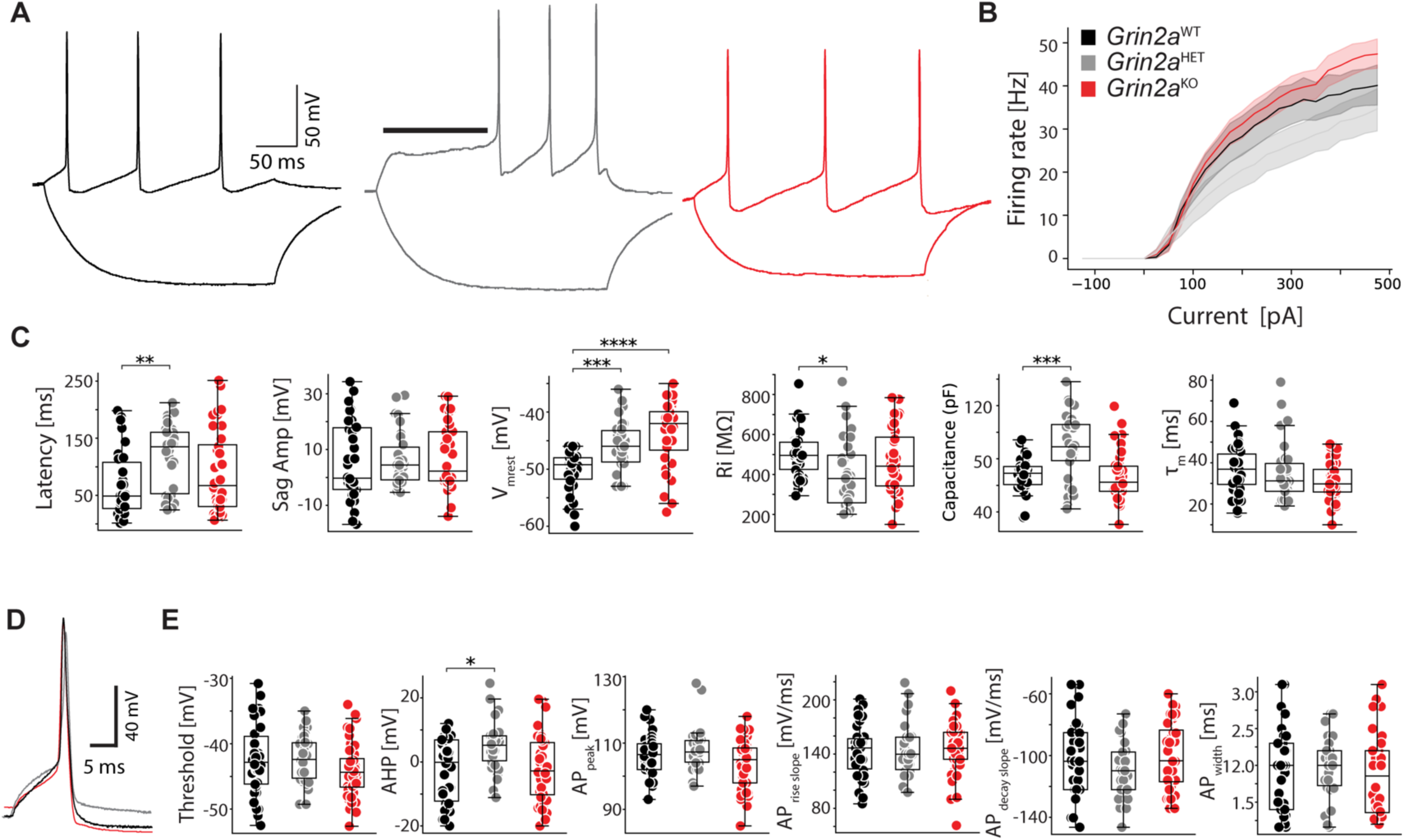
Intrinsic Active Properties of LC Neurons in *Grin2a* Mutant Mice. (**A**) Representative current-clamp recordings evoked repetitive firing of LC neurons from WT and *Grin2a* mutant mice in response to 300-ms current injections. (**B**) Input–output (I–O) curves plotting action potential firing rate versus current injection (−125 to +475 pA) across genotypes. (**C**) Quantification of intrinsic passive properties, including resting membrane potential (Vm), input resistance (R _in_), and membrane capacitance, in LC neurons from each genotype. (**D**) Representative action potential (AP) waveforms for WT and *Grin2a* mutant mice. (**E**) Quantification of AP waveform kinetics, including amplitude, threshold, rise/decay times, and afterhyperpolarization (AHP). Data are presented as mean ± SEM (*n* = 32– 40 cells from 6 mice per genotype). All statistical analyses were performed using the Kruskal–Walli’s test. Asterisks denote significance: * *p* < 0.05, ** *p* < 0.01, *** *p* <0.001, **** *p* <0.0001. Box plot centers represent medians, box edges the 25th and 75th percentiles, and whiskers the minimum and maximum values.

**Figure S2.**
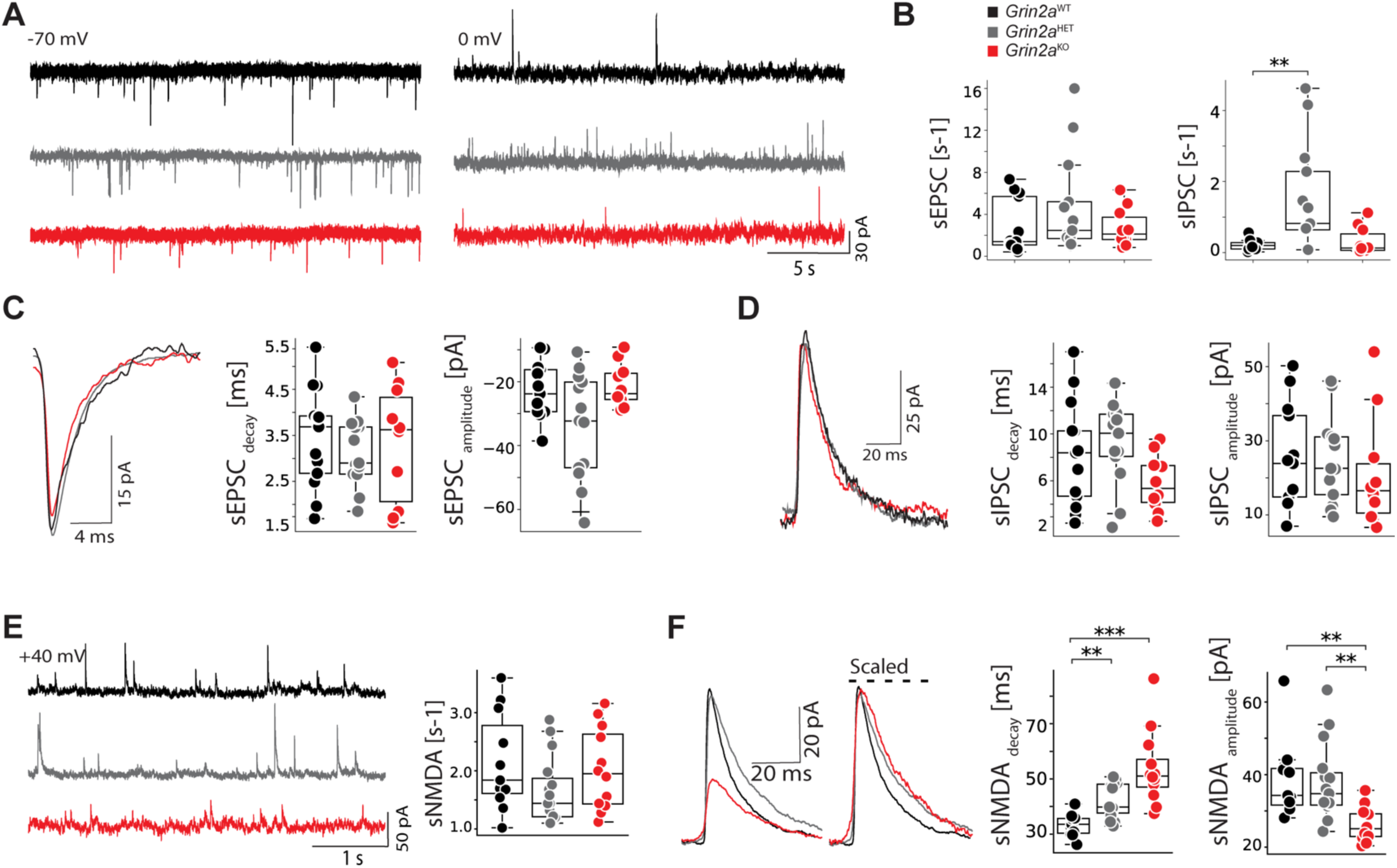
Spontaneous Excitatory, Inhibitory, and NMDA Currents in LC Neurons of *Grin2a* Mutant Mice. (**A**) Representative whole-cell patch-clamp recordings of spontaneous excitatory postsynaptic currents (sEPSCs) at −70 mV (left) and spontaneous inhibitory postsynaptic currents (sIPSCs) at 0 mV (right) in LC neurons from WT, *Grin2a ^HET^*, and *Grin2a ^KO^* mice. (**B**) Box plots comparing sEPSC and sIPSC frequencies across genotypes. **(C, D)** Representative waveforms (**C**) and box-plot quantifications (**D**) of sEPSC and sIPSC amplitudes and kinetics (e.g., rise and decay times). (**E**) Box plots of spontaneous NMDAR-mediated currents (sNMDA) at +40 mV. Traces on the left illustrate typical sNMDA events across genotypes. (**F**) Representative individual sNMDA currents (left) and box plots (right) summarizing sNMDA amplitude and decay time across genotypes. Data are presented as mean ± SEM (*n* = 11–14 cells from 3 mice per genotype). Statistical analysis was performed using the Kruskal–Walli’s test. Asterisks denote significance: * *p* < 0.05, ** *p* < 0.01, *** *p* <0.001. Box plot centers represent medians, box edges the 25th and 75th percentiles, and whiskers the minimum and maximum values.

